# Origin and Development of the Claustrum in Rhesus Macaque

**DOI:** 10.1101/2023.02.06.527286

**Authors:** Hong Li, Alvaro Duque, Pasko Rakic

**Author notes:** Corresponding author: Pasko Rakic, **Email:**. **Author Contributions:** HL conceived the idea, executed experiments, and wrote the paper; HL and PR designed the study; AD provided images from MacBrain Resource, and AD and PR edited the paper, and provided scientific advice and funding sources.

## Abstract

Understanding the claustrum’s functions has recently progressed thanks to new anatomical and behavioral studies in rodents, which suggest that it plays an important role in attention, salience detection, slow-wave generation, and neocortical network synchronization. Nevertheless, knowledge about the origin and development of the claustrum, especially in primates, is still limited. Here, we show that neurons of rhesus macaque claustrum primordium are generated between embryonic day E48 and E55 and express some neocortical molecular markers, such as NR4A2, SATB2, and SOX5. However, in the early stages, it lacks TBR1 expression, which separates it from other surrounding telencephalic structures. We also found that two waves of neurogenesis (E48 and E55) in the claustrum, corresponding to the birthdates of layers 6 and 5 of the insular cortex, establish a “core” and “shell” cytoarchitecture, which is potentially a basis for differential circuit formation and could influence information processing underlying higher cognitive functions of the claustrum. In addition, parvalbumin-positive interneurons are the dominant interneuron type in the claustrum in fetal macaque, and their maturation is independent of that in the overlaying neocortex. Finally, our study reveals that the claustrum is likely not a continuance of subplate neurons of the insular cortex, but an independent pallial region, suggesting its potentially unique role in cognitive control.

**Significance Statement:** The claustrum is believed to have a role in many high cognitive functions. However, the origin and development of this mysterious structure remain unknown, and the understanding of its relationship with the neocortex is ambiguous. Here we examined neuron origin and development of claustrum in rhesus macaque during the prenatal and postnatal periods. We found that the claustrum is formed as an independent telencephalic area as early as E55, and it seems not related to the subplate of the insula, although it shares some molecular characteristics with the neocortex. The claustrum excitatory neurons are generated sequentially around E48 and E55 and build a “core and shell” structure that may be a basic computing neuronal circuit unit underlying higher cognitive functions.

## Introduction

The claustrum is an elusive and mysterious structure comprising a thin sheet of cells sandwiched between the external capsule and the extreme capsule located below the insular cortex in primates. It is important for higher cognitive functions and it is involved in pathogenesis of psychiatric diseases. More than a decade ago, Crick and Koch (1) suggested that the claustrum plays a role in consciousness. Their integrated conscious percept model was based on the claustrum’s widespread connectivity profile, with identified projections to and from the sensory, motor, and associative cortical areas (2– 7). Knowledge of the physiological functions of the claustrum evolved in the past few decades as advanced anatomic and electrophysiological approaches were applied to claustrum studies in various species (7–15). The forefront discovery of claustrum physiology function is that the massive connectivity between the claustrum and neocortex enables the claustrum to synchronize neuronal activity between cortical areas (1, 16) as well as to generate and regulate slow waves during deep sleep, especially in the prefrontal and anterior cingulate cortices (14). The claustrum involves a broad range of general cognitive functions including attention, resilience to distraction, and cognitive action control (7, 10, 11, 15, 17). Furthermore, there is increasing evidence from human studies that the claustrum plays a role in psychiatric diseases. For example, reduced volume and gray matter of the bilateral claustrum, are found in postmortem patients with schizophrenia and major depressive disorder (18, 19). Functional magnetic resonance imaging (fMRI) studies in schizophrenia patients found that the intensity of delusions and hallucination is inversely related to the volume of the left claustrum and right insula, but not cerebral gray matter volume, suggesting that the claustrum may be a potential center for delusions and hallucinations (20).

Given the important roles of the claustrum in higher cognitive functions and potential neuronal mechanisms underlying psychiatric diseases in humans, it is pivotal to understand where and when the claustrum cells originate and how the claustrum develops in the primate brain. Though the claustrum exists in almost all vertebrate brains, in species other than primates it lacks homology with humans since the extreme capsule separating the claustrum from the insula cortex is only present in primates (21, 22). The endopiriform nucleus, a ventral adjacent nucleus in rodents, does not exist in primates (23). Historically, the claustrum origin and development have gone through different theories of belonging to basal ganglia, being independent or being a layer of the insular cortex. As of now, molecular biology and transcriptomic approaches identified a group of genes that are expressed in the claustrum, the subplate and deep layers of the insula (21, 24). It is suggested that the claustrum may share its origin with the insula from lateral pallium (LPALL) (22, 24). However, it is perplexing that *in utero* electroporation of LPALL with EGFP at the time of subplate neurogenesis labeled primarily the insular cortex, with no apparent claustrum labeling (24). The above studies focus on rodents and developmental data from primate brains are still lacking. In the present study, we systematically investigated the development of the rhesus macaque claustrum from early embryonic stages to adult. We demonstrate that the claustrum of rhesus macaque exhibits unique molecular characteristics at a very early stage and during development and that it shares some gene expression with the neocortex early on and along its developmental trajectory. We pinpoint the time of origin of the neurons in the claustrum with a combination of analysis using archived tritiated thymidine birth-dating experiments from Collection 1 of the MacBrain Resource and modern immunofluorescence dual birth-dating methods. Our results suggest that the claustrum is an independent pallium region that has a unique origin different from that of the subplate of the neocortex.

## Results

### Principal neurons in the “core” were born around E48

To determine when claustrum neurogenesis occurs in relation to the cortical neurons, we used the same DNA labeling method as in a previous publication (25). Thus, we analyzed tritiated thymidine injections on timed pregnant rhesus macaque experiments available in MacBrainResource Collection 1 (https://medicine.yale.edu/neuroscience/macbrain/). Adult and postnatal macaque brain sections with tritiated thymidine injections at E45 to E75 were screened for labeling in the claustrum. We found several cases of positive staining in the ventral claustrum but not the middle and dorsal parts with E47-E50 tritiated thymidine injections (**Fig. 1A-C**). To further investigate in detail the birthdates of claustrum neurons, we injected CldU (5-Chloro-2’-deoxyuridine) at E40 and IdU (5-Iodo-2’-deoxyuridine) at E46, in the same pregnant dam, and sacrificed at E73. This approach using multiplex immunofluorescence helped reduce the number of animals needed and allowed immediate comparison of cells born at two different ages. As shown in (**Fig 1D, E**), neurons born at E40 (CldU) are not present in the claustrum, but abundantly in surrounding areas including the subplate of the insular cortex, lateral migratory stream (LMS) as well as ventral pallial structures (**Fig. 1D**). Similarly, neurons born at E46 (IdU) are not found in the claustrum, but instead, are abundant in the cortical plate of the insula (**Fig. 1E**).

**Fig. 1.**
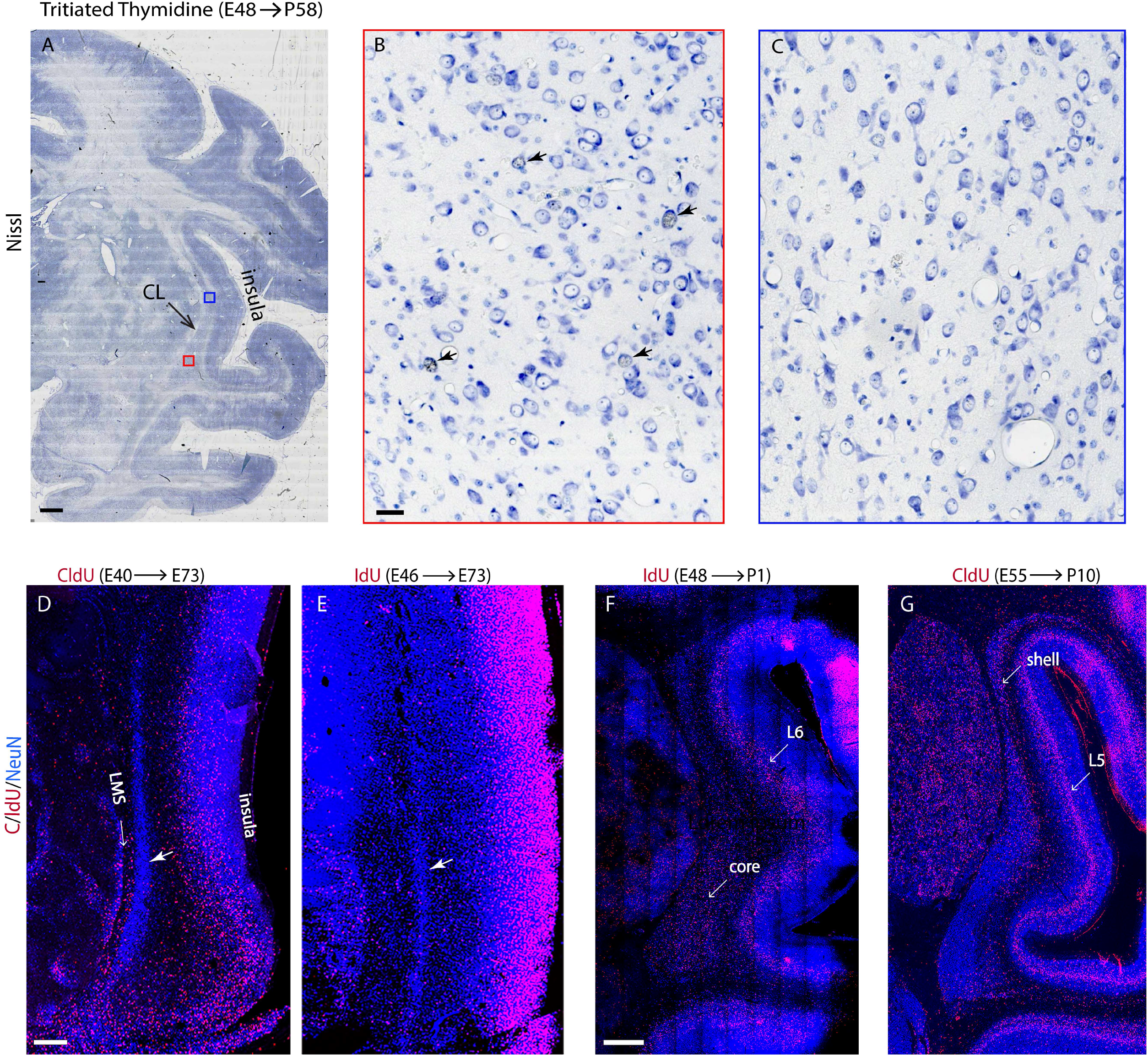
Claustrum neurons were born at E48 and E55. (**A-C**) Representative nissl staining images of tritiated (H_3_) thymidine birth-dating experiment from MacBrain Resources (Collection 1) demonstrated that the “core” neurons but not the “shell” neurons in claustrum were labeled with tritiated (H_3_) thymidine when injected on E48. (**B**) and (**C**) are enlarged images from color-coded framed areas in (**A**). Arrowheads in (**B**) point to 3 neurons labeled with H_3_-thymidine in the “core”. (**D-G**) A summary of series of multiplex deoxyuridine (CldU and IdU) injection experiments performed from E40 to E70. with either double IdU and CldU injections (**D** and **E**), or single IdU (**F**) or CldU (**G**) injections. White arrow heads in (**D)** and (**E**) point to claustrum. LMS: lateral migratory stream; L6: neocortex layer 6; L5: neocortex layer 5; Scale bar: A: 2mm; B, C: 30μm, D, E: 500µm; F, G: 1mm.

In a different pregnant dam, ldU was injected at E48 and sacrificed at postnatal day 1 (P1). Neurons born at E48 were found in the ventral part of the claustrum (**Fig. 1F, 2A-C)** and observation of serial sections anterior to posterior revealed that those neurons are present in sections in the ventral but not in the medial and dorsal and ventralmost of the claustrum (**Fig. 2A-D**). Our results demonstrated that the first wave of neurogenesis in the claustrum occurred by E48 forming a “core” (**Fig. 1F, 2D)**. In the neocortex, layer 6 neurons of the insula were born at this age, but barely subplate neurons are detected (**Fig. 1F**).

**Fig. 2.**
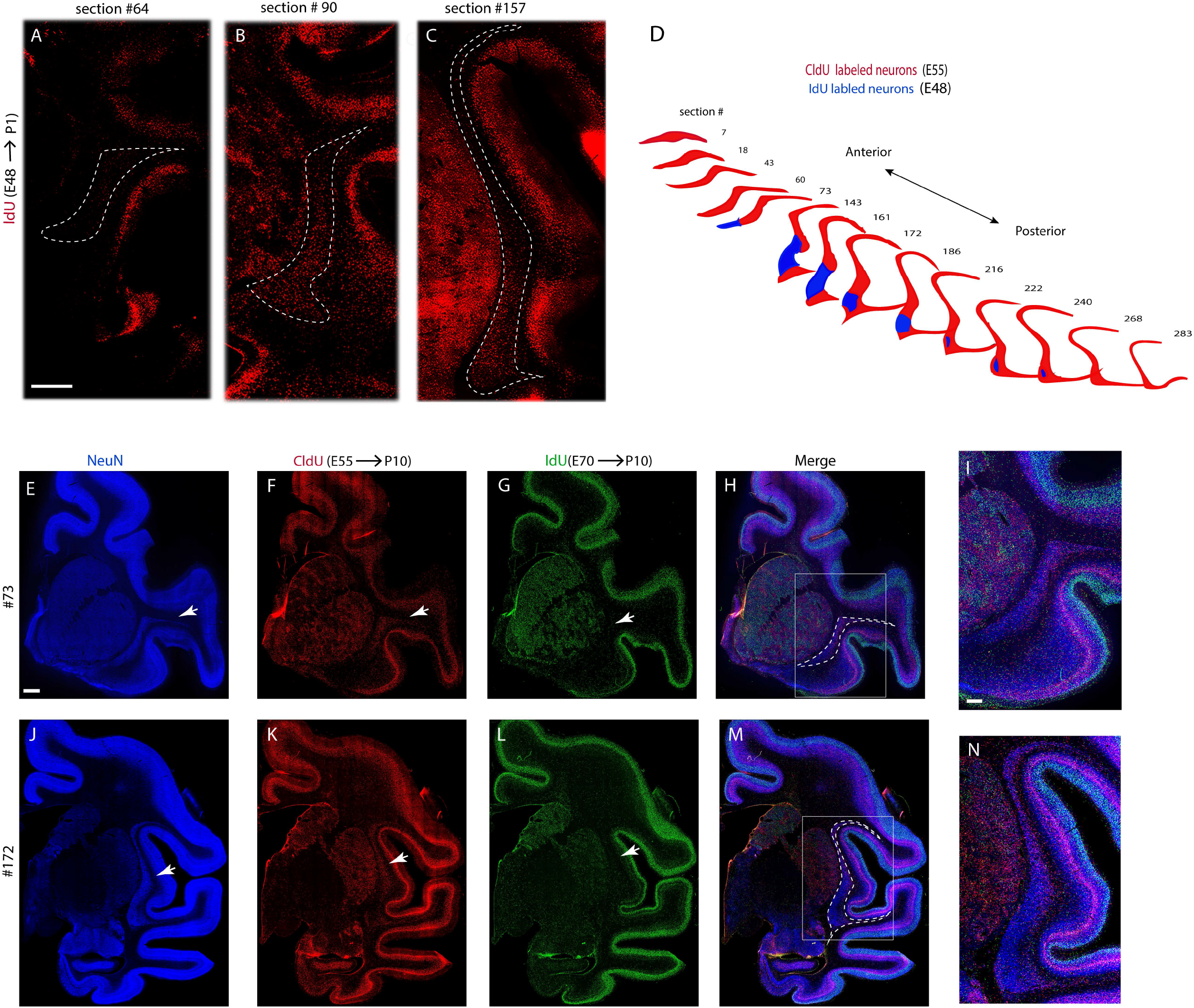
Claustrum neurons birth-dating experiments performed with CldU and IdU delineate a “core” and “shell” structure. (**A-C**) A series of anterior to posterior sections show claustrum “core” neurons were labeled when IdU was injected at E48. Dashed lines are outline of the claustrum. (**D**) A cartoon 3D model of the two waves of claustrum neurogenesis. The “core” is represented in blue color, and the “shell” is represented in red. (**E-N**) A dual injection of CldU and IdU performed at E55 (CldU) and E70 (IdU) shows that neurogenesis of the claustrum “shell” occurs at E55 (arrowheads). Dashed lines in (**H**) and (**M**) are outlines of the claustrum. (**I**) and (**N**) are enlarged images of white framed areas in (**H**) and (**M**) respectively. Scale bar: A-C: 2mm; E-H, J-M: 2mm; M, R: 1mm.

### Principal neurons in the “shell” were born by E55

We performed another multiplex CldU/IdU injections at ages E55, and E70, and discovered neurons in the medial to dorsal regions of the claustrum and the ventralmost thin sheet as well as anterior and posterior ends of the claustrum neurons were born by E55 (**Fig. 1G, 2D-N**). We described those neurons together as the “shell” of the claustrum. In this temporal window, insula cortical layer 5 neurons were born as demonstrated by co-staining of CldU and NeuN (**Fig. 1G**). There is no claustrum neuron labeled with IdU injected at E70 (**Fig. 2G, L**). In general, 3D reconstruction of the two waves (E48 and E55) of neurogenesis divides the claustrum into two compartments: “core” and “shell” (**Fig. 1F-G**). These results are summarized in **Fig. 2D**.

### Claustrum primordium can be detected molecularly as early as E55 in the embryonic brain of rhesus macaque

To determine molecular properties of the claustrum primordium and to compare them with neocortex during embryonic development of rhesus macaque, we examined the immunoreactivity of NR4A2 as early as E55. We found a cluster of postmitotic neurons that exhibit strong expression of NR4A2 deep in the insular cortex at this stage (**Fig. 3A**). This is compared to very low or no expression of NR4A2 detected in the neocortex at E55 (**Fig. 3A**). As the embryonic brain develops, the NR4A2 positive claustrum primordium elongates into a long dorsal to ventral strip, as seen in coronal sections (**Fig. 3B-E**). At and after E60, weak and scattered expression of NR4A2 is found in deep layers of the neocortex while the expression in the claustrum remains prominent until adulthood (**Fig. 3B-E**). SATB2 is expressed in the claustrum primordium and cortical plate of the lateral neocortex at E55 as well. Its expression in the claustrum is maintained throughout embryonic development into adulthood while the expression in the neocortex follows the maturation of cortical neurons from lateral to medial during development (**Fig. 3F-O**). NeuN was not detectable at E55 but was highly expressed in the claustrum primordium starting as early as E60 (**Fig. 4A-E**) indicating that claustrum neurons are early to mature. SOX5, a transcription factor that is expressed in neocortical layers 5 and 6, and the subplate in rodents (26), is highly expressed in the claustrum primordium at E60 as well as layers 5, 6, and the subplate of the neocortex in embryonic macaque brains (**Fig. 4F, G**), consistent with the previous study in rodents. These results demonstrate that the developing claustrum has some overlapping expression profiles with the neocortex but that many of the same neocortical markers are uniquely expressed or expressed at earlier timepoints in the claustrum, suggesting that the claustrum deviation from the neocortex may occur at the progenitor level.

**Fig. 3:**
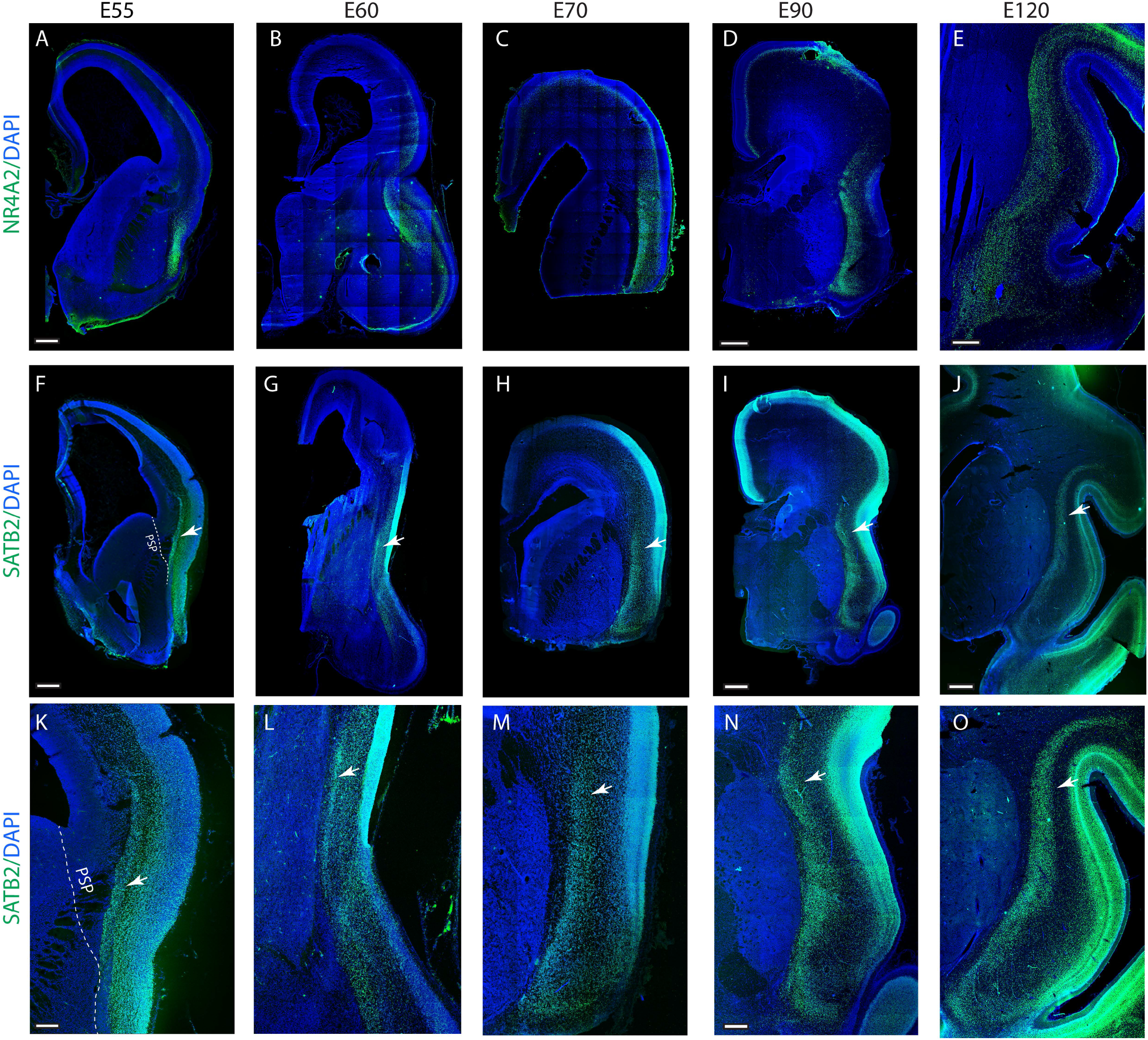
NR4A2, SATB2 immunostaining mark earliest primordium of macaque claustrum. (**A-E**) NR4A2 protein expression labels early claustrum primordium at E55 and maintains the specific expression throughout embryonic and fetal stages. (**F**-**J**) SATB2 protein is detected in the claustrum (arrowheads) earliest at E55 and the expression is high throughout fetal development. (**K**-**O**) are enlarged images from (**F**-**J**). PSP: pallium and subpallium borderline. Scale bar: A-C, E, O: 2mm; F-H: 1mm; D, I: 1.5mm; J: 2mm; K-M: 250µm; N: 500µm.

**Fig. 4:**
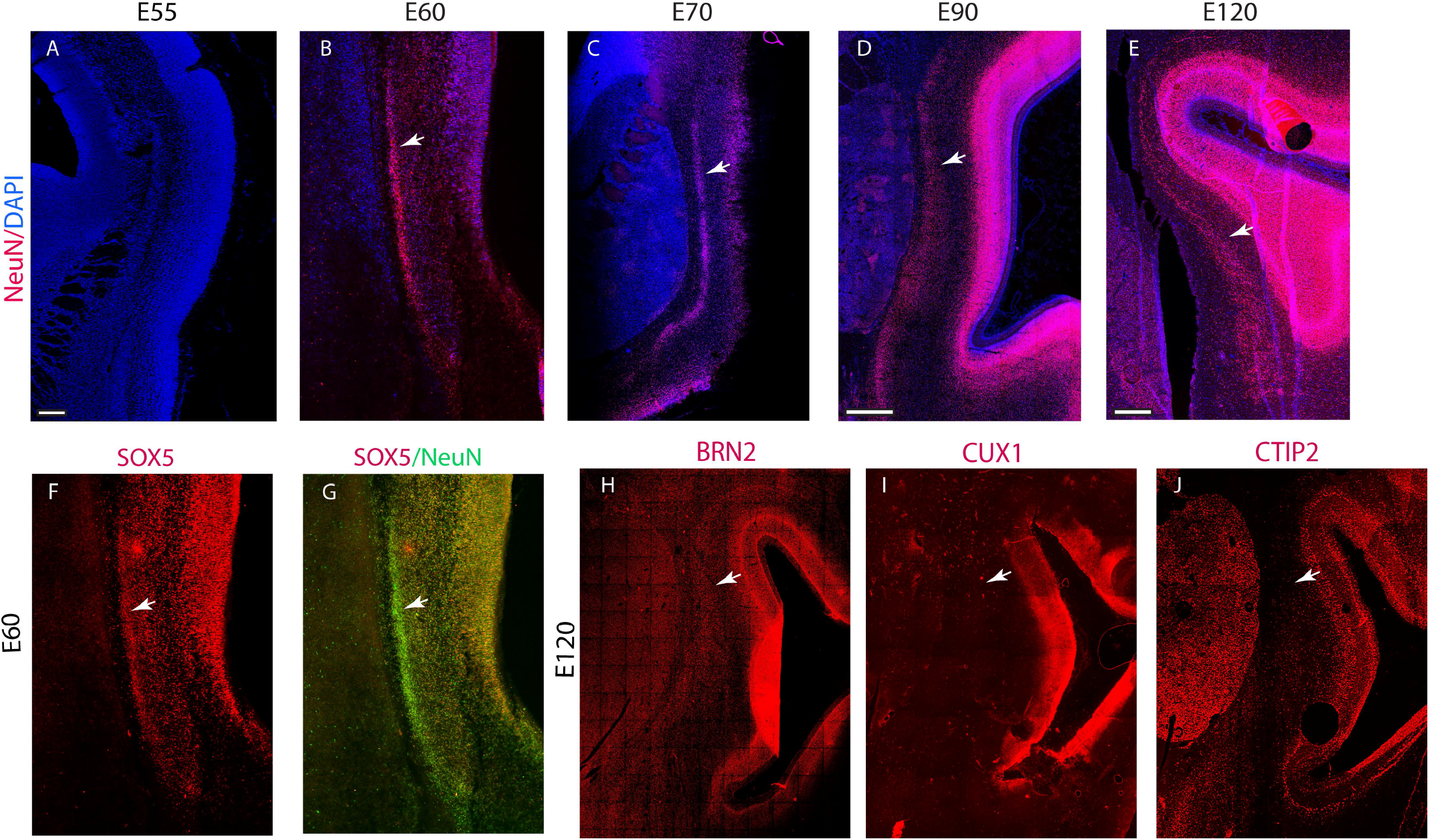
Molecular characteristics of the developing claustrum. (**A**-**E**) NeuN is expressed in the claustrum primordium and the neocortex as early as E60 and is highly expressed throughout fetal development. White arrow heads point to the claustrum. (**F, G**) SOX5 is expressed in the claustrum (arrowheads) at E60. (**C**) BRN2 is detected at E120 in the claustrum, but CUX1 and CTIP2 are absent. White arrow heads point to the claustrum. Scale bar: A-C, F, G: 250 µm; D: 1.5mm; E, H-J: 1mm.

We also examined the expressions of other neocortical neuronal markers, such as CUX1, BRN2, CTIP2, FOXP2, and TBR1 in embryonic and fetal developmental stages (**Fig. 4H-J**). We found the prenatal claustrum of the macaque expresses the upper layer marker, BRN2, and the expression starts around E70 and peaks at about E120 (**Fig. 4H**), but down-regulates to an undetectable level after birth. The expression of BRN2 is higher in the dorsal claustrum than in the ventral part of the claustrum. CUX1, CTIP2, (**Fig. 4I, J**) and FOXP2 (Data not shown) are absent in embryonic brains. Most importantly, TBR1 has a unique expression pattern in the embryonic claustrum (see details below).

### Claustrum excitatory neurons are organized into a TBR1 negative “core” and a TBR1 positive “shell” pattern at embryonic and fetal stages

The TBR1 expression in the prenatal claustrum is profoundly different from other molecules we examined. At E55 and E60, TBR1 is not expressed or expressed at very low levels in the claustrum primordium in comparison to its surrounding pallial neurons. In addition, the claustrum primordium can be recognized with a low expression hollow located near the pallium and subpallium border in coronal sections with TBR1 immunostaining (**Fig. 5A, B, F, G**). As the claustrum develops, the differential expression of TBR1 within the claustrum becomes even more blatant. At E70, cells in the circumference of the claustrum primordium express TBR1 moderately, forming a thin sheet of TBR1 positive cell “wall” which encircles TBR1 negative or low expression neurons but SatB2/NeuN positive neurons (**Fig. 5C, H, D, I**). The notable differential expression of TBR1 within the claustrum continuously presents at late fetal stages. By E120, when the claustrum is further separated from the neocortex, the claustrum displays a distinctive TBR1 negative “core” and TBR1 positive “shell” structure demonstrated by co-immunostaining with NeuN (**Fig. 5E, J**). The TBR1 positive neurons form a thin sheet wrapping around the “core” of the claustrum primordium and elongate dorsally and laterally towards the temporal lobe as the claustrum develops. The general “core” and “shell” cytoarchitecture of TBR1 distribution becomes less obvious after birth. TBR1 negative neurons and positive neurons were found intermixed throughout the claustrum of postnatal brains. The above results show that claustrum principal neurons and neocortical excitatory neurons share many molecular features, but differential expression of TBR1 within the fetal claustrum sets the trajectory of claustrum development apart from that of the neocortex.

**Fig. 5:**
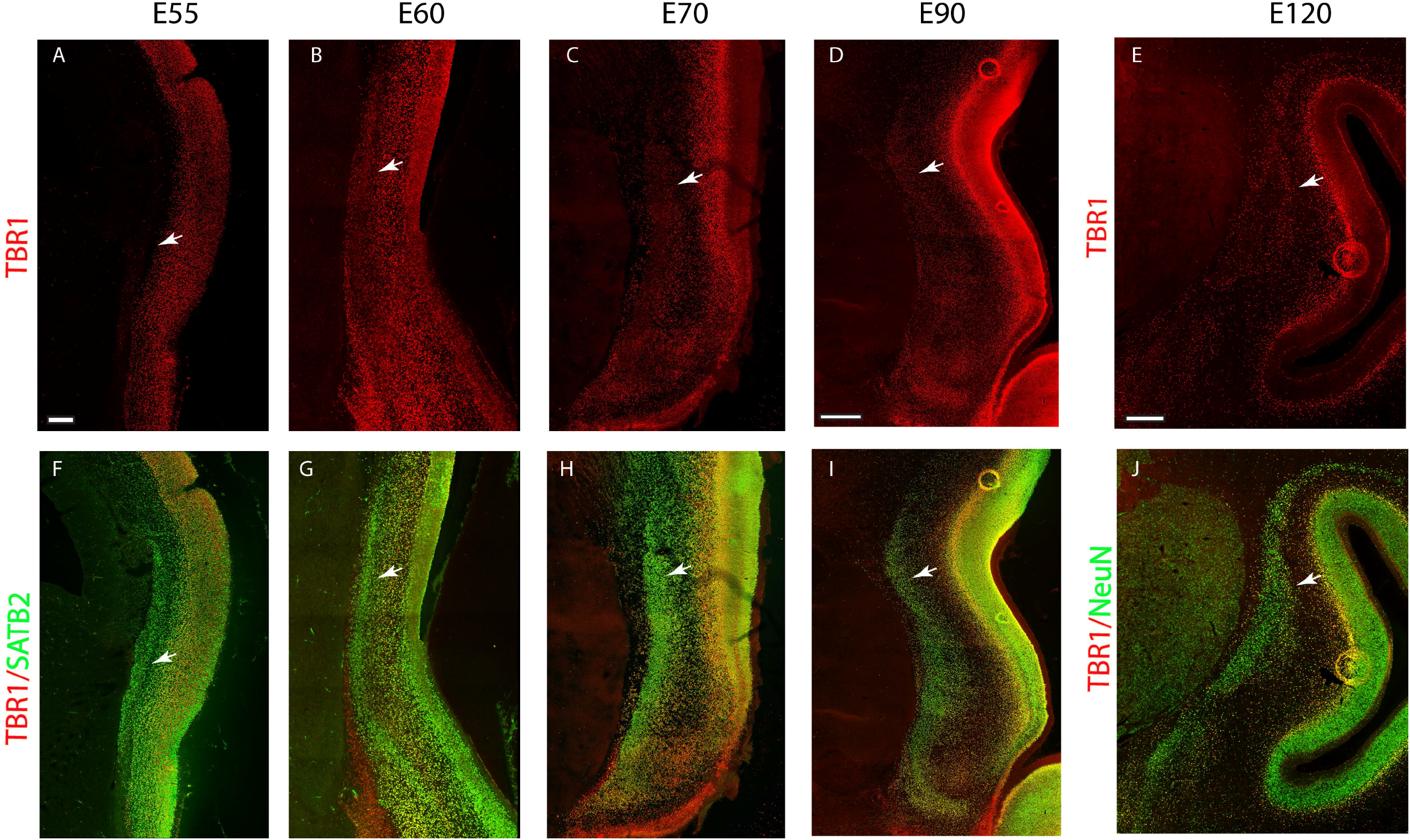
Absence of (or low) TBR1 expression in the claustrum primordium at E55 and E60, and the “core” region of the claustrum at mid-late fetal stages. (**A, B**) TBR1 protein expression is much lower in the claustrum (arrowheads) than adjacent regions from E55 to E60. (**C**-**E**) As claustrum develops, TBR1 positive “shell” encircling TBR1 negative “core” is present from E70-E120. (**F-J**) SATB2 or NeuN coimmunostained with TBR1 shows that neurons in the “core” of the claustrum express no or very low TBR1. Scale bar: A-C, F-H: 250 µm; D, I: 1.5mm; E, J: 1mm.

### Neurogenesis and molecular phenotype of the “core” and “shell”

We then asked whether neurons in the “core” and “shell” of the claustrum according to their birth dates are different from the TBR1 immunoreactivity “core” and “shell”. Thus, we examined the level of colocalizations of TBR1 with IdU (E48) and CldU (E55) as well as SATB2 with IdU (E48) and CldU (E55). Unlike TBR1, SATB2 is uniformly expressed in the “core” and “shell” neurons throughout the developmental stages investigated (**Fig. 3F-O**). Double immunostaining of IdU (E48) with TBR1 and SATB2 indeed showed that there are fewer TBR1/IdU colocalized cells (**Fig. 6A-C, M, O**) than SATB2/IdU colocalized cells (**Fig. 6D-F, N, O**) in the claustrum, indicating that the majority of TBR1 negative neurons in the claustrum are born at ∼E48. Colocalization analysis of CldU (E55) with TBR1 and SATB2 did not show much difference between TBR1/CldU (**Fig. 6G-I, P, R**) and SATB2/CldU (**Fig. 6J-L, Q, R**), indicating TBR1 positive neurons are born at ∼E55. However, these molecular difference between the “core” and “shell” is prominent only during prenatal development, especially in the late fetal stages. The differential molecular expression between “core” and “shell” cytoarchitecture within the claustrum is not distinguishable in late postnatal stages and adults. Together, these cytoarchitectural experiments demonstrate the claustrum is formed independently from the neocortex.

**Fig. 6:**
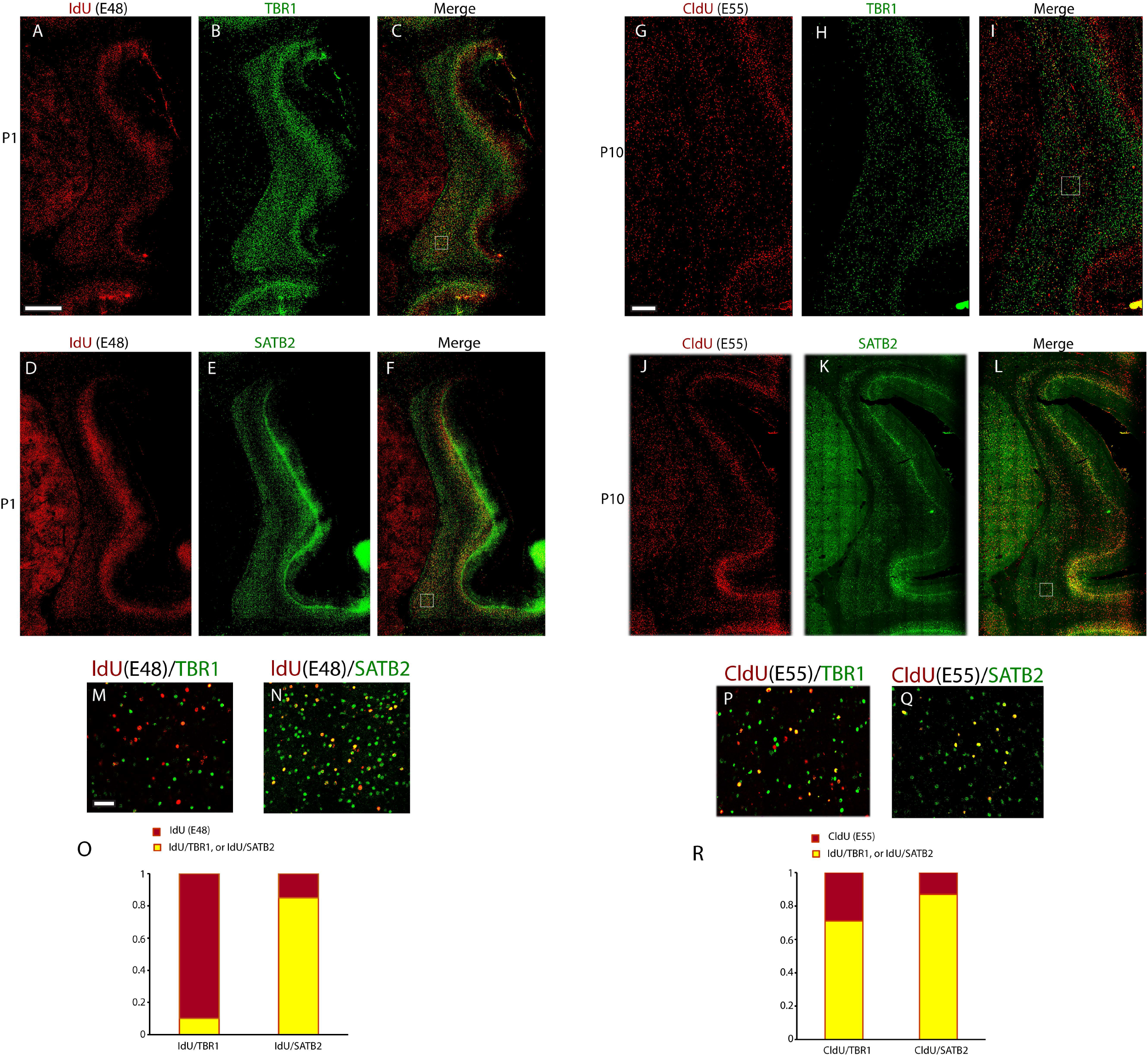
Neurogenesis and molecular property of the “core” and “shell”. (**A-F**) Low magnification images of the claustrum co-immunostained with CldU (injected at E48) and either TBR1 (**B, C**) or SATB2 (**E, F**). (**M**) and (**N**): High magnification images of the co-immunostaining in boxed areas in (**C**) and (**F**) respectively. (**O**) A histogram shows the percentage of CldU/TBR1 and CldU/SATB2 colocalized neurons to total CldU per unit. (**G-L**) Low magnification images of the claustrum with co-immunostained with ldU (injected at E55) and either TBR1 (**H-I**) or SATB2 (**K-L**). (**P**) and (**Q**): High magnification images of the co-immunostaining in boxed areas in (**I**) and (**L**), respectively. (**R**) A histogram shows the percentage of IdU/TBR1 and IdU/SATB2 colocalized neurons to total IdU per unit. Scale bar: A-F, J-L: 1mm; G-I: 500µm; M, N, P, Q: 50µm.

### Postnatal claustrum molecular features in rhesus macaque

We then investigated the molecular characteristics of the claustrum in postnatal rhesus macaques. SATB2 (**Fig. 7A, F, K**) and NeuN (**Fig. 7B, G, L**) are strongly expressed in the claustrum as well as in the neocortex at postnatal ages (from P7 to 1 year old) similar to that at embryonic stages. Expression of TBR1 in the claustrum after birth starts to lose its differential “core” and “shell” pattern, appears grossly even distributed (**Fig. 7C, H, M**). We further examined the protein expression of LATEXIN, an endogenous carboxypeptidase inhibitor previously found to be expressed in cat and rat claustrum (27, 28). Here, we found that in early postnatal ages (e.g., P7 and up to one year) LATEXIN is highly expressed in the claustrum but rarely present in the insula and striatum (**Fig. 7D, I, N**). By examining other brain areas in a series of coronal sections from P7 to adults, we observed that LATEXIN is also strongly expressed in entorhinal cortex, subiculum and hippocampus, and moderately in layer 2, 6 of the anterior cingulate cortex (ACC) (**Fig. 7D**). Furthermore, NR4A2 (NURR1) protein is expressed distinguishably in the claustrum of the postnatal macaque brains, but not in surrounding structures except some neocortical layer 6 neurons (**Fig. 7E, J, O**). These data demonstrate that the primate claustrum not only expresses neocortical excitatory molecular markers but also exhibits gene expression patterns that are unique and specific as compared to the surrounding structures.

**Fig. 7:**
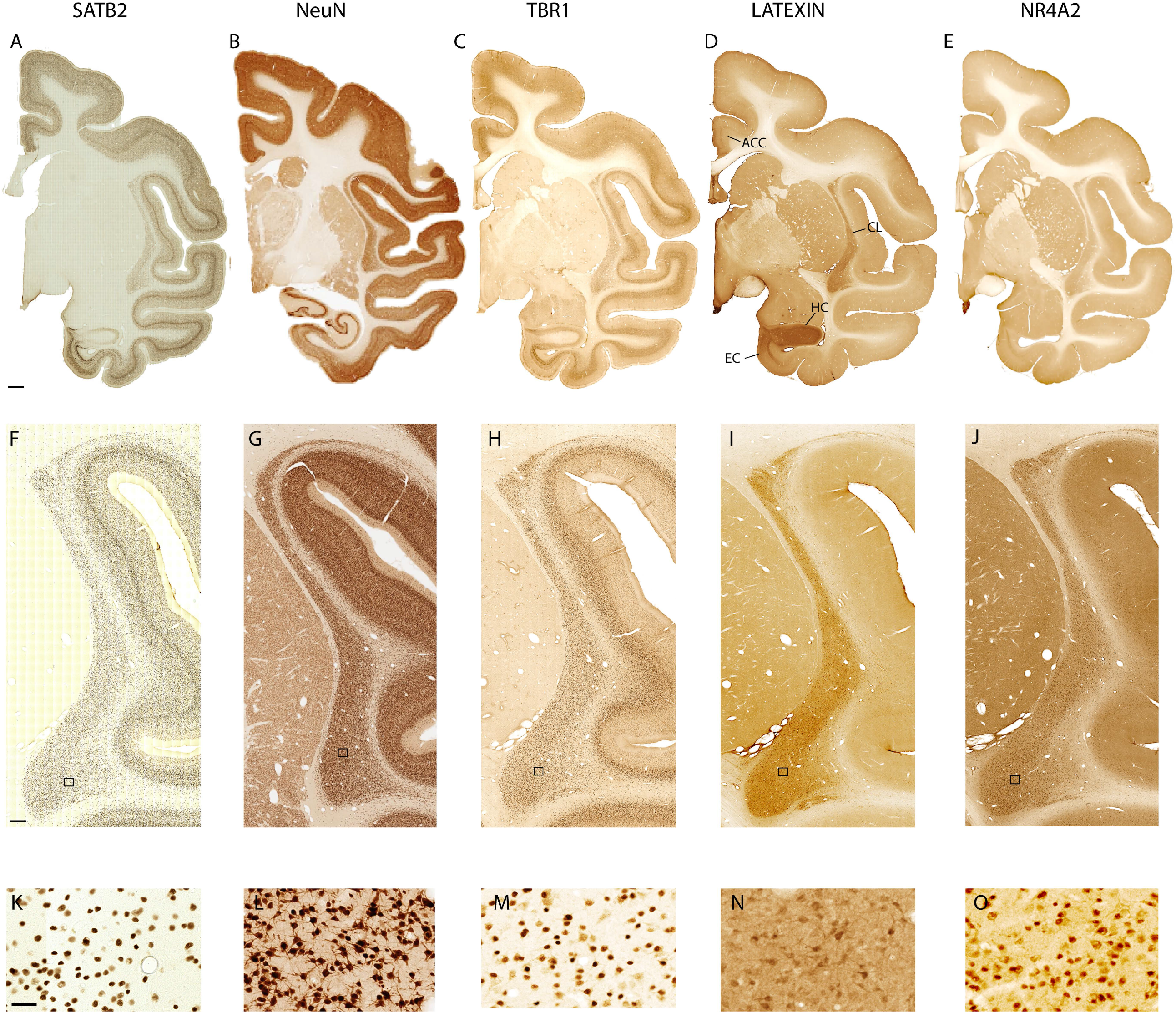
Claustrum expresses typical neocortical molecular markers as well as unique genes in postnatal rhesus macaques. Expressions of SATB2 (**A, F, K**), NeuN (**B, G, L**,), TBR1 (**C, H, M**), Latexin (**D, I, N**) and NR4A2 (**E, J, O**) are detected in the claustrum of postnatal rhesus macaque brains. SATB2, TBR1, Latexin, and NR4A2 are from a 1-year-old brain. NeuN immunostaining is from a P7 brain. **K, L, M, N, O** are close-up images in framed areas in **F, G, H, I, J**, respectively. ACC: anterior cingulate cortex; CL: claustrum; HC: hippocampus; EC: entorhinal cortex. Scale bar: A, C, D, E: 2mm, F, H, I, J: 500µm, K, M, N, O: 50µm. (MacBrainResource Collection 6).

### The development of interneurons in the claustrum

Previous studies showed that interneurons are abundant in the claustrum of rodents (29, 30). Here we explored what types of interneurons are in the developing claustrum of the rhesus macaque, when they emerge and whether their development is correlated with that in the neocortex or striatum. In the neocortex, there are three main types of interneurons, which account for almost all interneurons in the neocortex, molecularly marked by parvalbumin, somatostatin (SST), and 5-HT3aR (31). We immunostained parvalbumin, SST, and 5-HT3aR on E70, E90, E120, P1, and P10 macaque brains. There is no detectable signal in 5-HT3aR immunostained sections at embryonic and fetal stages, but we observed weak signals of parvalbumin immunostaining in the claustrum at E90, those neurons distribute evenly throughout the claustrum in fetal brains. In contrast, abundant parvalbumin positive neurons migrating to the neocortex are found accumulating in the subplate and deep layers of the neocortex. At E120, the immunoreactive signal from parvalbumin positive interneurons in the claustrum increases dramatically and evenly distributes throughout the claustrum (**Fig. S1A, B**). We observed that parvalbumin positive neurons in the claustrum have reached their permanent positions whereas those in the neocortex are still migrating, indicating the development of parvalbumin positive interneurons in the claustrum has its own trajectory. The expression is maintained in the claustrum postnatally and in adults (**Fig. S1C-H**). SST immunoreactivity is detected in the claustrum at a late fetal stage, E143 (**Fig. S1I, J**), and the signal increases slightly in the newborn and maintains up to adult (**Fig. S1K-P**). The density of SST positive neurons in the claustrum is similar to the deep layers of the neocortex throughout development (**Fig. S1I, K-P**). Other subsets of interneurons we analyzed are calbindin, calretinin, and NPY positive interneurons. Calbindin and calretinin positive neurons are not detectable during embryonic and fetal stages but are present in the claustrum after birth to adult (**Fig. S2A-L**). The density of these neurons in the claustrum also resembles that in the deep layers of the neocortex (**Fig. S2B, E, H, K**). Importantly and interestingly, NPY expressing neurons are low or absent in the claustrum but abundant in neocortex deep layers and striatum throughout macaque development (**Fig. S2M-P**). We did not find differential distribution between the “core” and “shell” in any subtypes of interneurons examined.

## Discussion

### Neurogenesis of claustrum principal neurons in primates

We found that there are two major waves of neurogenesis in the making of the claustrum in the macaque developing brain. Neurogenesis of the macaque claustrum does not occur during the same temporal window as the subplate neurons of the insula; but rather at the time when neocortical layer 5 and layer 6 excitatory neurons were born. A series of deoxyuridine injections from E40 to E70 (**Fig.1 & Fig. 2**) demonstrated that claustral neurons in the macaque were born after the subplate of the insula cortex. Instead of an “inside-out” model for neocortical laminar neurogenesis, the claustrum’s “core” is formed first and then followed by a “shell”, which wraps around the “core” while extending dorsally and laterally, forming a unique cytoarchitecture reminiscent of a brain nucleus. A few rodent birth-dating experiments studies yielded inconsistent results. For example, autoradiographic labeling (H^3^-thymidine) studies performed on the mouse at early embryonic stages found mouse claustrum labeling as of E11, and finished neurogenesis at E13 (32, 33). Similar experiments performed on rats observed that claustral populations are produced between E11.5 and E14.5 with a peak at E12.5 (22). The most recent study by Hoerder-Suabedissen (34) on neurogenesis of the claustrum in mice indicated that claustrum neurons were born at a short time window, E12.5 and neurons in endopiriform nucleus were born at E11.5. Here we found that claustrum neurons in macaques are generated later than subplate neurons of the insular cortex. Hence, formation of the claustrum is an independent process involving a separate set of neuronal progenitors which may diverge from the neocortical progenitors early on. To discover the progenitors in the ventricular sites where claustral neurons are produced, identifying a molecular marker specific to the progenitors is needed.

### The “core” and “shell” cytoarchitecture of the developing claustrum

We revealed “core” and “shell” structures in the claustrum of rhesus macaque in two different ways, one with TBR1 immunostaining and the other with birth-dating experiments. These results support Ramon y Cajal’s (1902) suggestion that the claustrum was a pallial nucleus rather than a cortical layer (22). Core and shell structure in adult claustrum has been suggested in mice and monkeys in some studies with different perspectives, for example, visual and anterior cingulate cortical inputs, parvalbumin positive interneurons and their neurites are distributed in the core whereas auditory and orbitofrontal cortical inputs terminate in the peripheral shell (9, 21). Binks et al (2019) (21) described that *Nr4a2, Ntng2*, and *Synpr* are expressed in the core and the expression of *Crym* is in the shell in both adult rodents and macaques. Here, in developing macaque claustrum, the “core” is the enlarged ventral part of the claustrum, and the “shell” is a thin sheet of cells warping around the “core” along with medial, dorsal and lateral extend of the claustrum. It is reasonable to conjecture that molecular characteristics, connectivity, and functions may be significantly different between the “core” and “shell”, and that a potential hierarchy for computation and information processing may exist between the two regions. Interestingly, the claustrum “core” and “shell” in macaque brains are detectable with TBR immunoactivity only during embryonic and fetal stages. After birth, differential expression of TBR1 between principle neurons diminished, and the “core” and the “shell” become indiscernible because TBR1 negative neurons start to catch up with their TBR1 expression. Compartmentalization and functional segregation in the claustrum have been reported in adult rodents (7, 8, 35) and macaques (36). A topographic map is formed with multisensory modalities in the dorsal and connectivity with the prefrontal cortex is in the ventral part of the claustrum. A study suggested that ventral claustrum neurons connect more densely with the neocortex than the dorsal claustrum (37). The differences in neurogenesis and molecular property within the claustrum during fetal development described here may underlie the functional segregation in mature claustrum in primates.

### The claustrum in rhesus macaque is an independent pallial nucleus rather than the subplate of the insula

Our study shows that the claustrum shares many gene expression features with the neocortex, but also revealed its unique molecular features and temporal control of neurogenesis that are different from the neocortex during early development. The expressions of SATB2 and NeuN in the claustrum at early embryonic stages demonstrates that claustrum principle neurons are telencephalic neurons which are generated early. Some neocortical laminar molecular markers detected in the claustrum were characterized by a mixture expression of upper- and lower-neocortical layer markers, for example, SOX5 and BRN2. Claustrum principle neurons are reported to be Emx1 lineage neurons in rodents (38). These findings seem to support that the claustrum is part of the subplate neuron pool of the neocortex. However, TBR1 absence in the claustrum primordium is surprising finding, and it sets the claustrum apart from all other pallium derivatives including the insula. It suggested that the claustrum excitatory neurons may derive from unique and different progenitors from the neocortical lineage. Therefore, the claustrum neurons originate in the pallium, but independently from the insula and adjacent areas of the neocortex.

### Interneurons in the macaque claustrum

Unlike principle neurons, interneurons in macaque claustrum distribute evenly throughout development and in adults. Interneurons in the claustrum and neocortex may come from the same progenitors in the ganglionic eminence. It is interesting to note that NPY expression are much lower in the claustrum than adjacent subplate of neocortex and striatum throughout development. We have not observed interneurons compartmentalization within the claustrum if expressed during development and in adults, consistent with a previous study (40). In rodents, parvalbumin and calbindin positive neurons are more abundant in the dorsal claustrum than in the ventral, usually referred to as the endopiriform nucleus (41).

### Species difference in the claustrum

The claustrum in mammalian species other than primates (macaques and humans) has different shapes. The shape of the claustrum in cats, ferrets, and pigs seems inverted as compared to the shape in macaques, the enlarged part is in the dorsal in coronal sections rather than in the ventral, called a “superior pyramidoid puddle” (42, 43). Whether the “superior pyramidoid puddle” is equivalent to the “core” of macaque defined in this study is not known. In rodents, claustrum (CLA) and endopiriform nucleus (En) often come together in most studies focusing on the claustrum. Analogue of endopiriform nucleus may not exist in macaques and humans. Only in marmosets, the CLA-En complex is described together as a cell-dense nucleus with strong NR4A2 mRNA signals (23). It is not known if a homologous endopiriform nucleus in macaques merges into the claustrum, becomes the ventral claustrum or if it disappeared during evolution. Traditional as well as viral tracing studies in rodents (9, 10, 44) and primates (45), showed similarity in anatomical connectivity between the endopiriform nucleus in rodents and ventral claustrum in macaques. In rodents, visual and auditory cortical areas, as well as prefrontal and olfactory areas, connect with the endopiriform nucleus, whereas in primates, the prefrontal cortex including dorsolateral and infralimbic areas as well as visual and auditory cortex connect with the ventral claustrum, (the “core” described in this study). In the domestic cat, the visual claustrum is in the “superior pyramidoid puddle” (42). The dorsal and medial “shell” of the primate claustrum connects with somatosensory and motor zones. It is worth noting that, in humans, the ventral claustrum (the “core”) is proportionally much larger than any other species in consistent with its enlarged prefrontal and visual cortical areas (23). It must be evolutionally required for its complex information processing and integration.

In conclusion, we provided evidence revealing the origin and development of the claustrum throughout the embryonic to fetal stages in the rhesus macaque. Our data support that the claustrum is a developmentally and cytologically independent telencephalic region that shares more molecular features with the neocortex than with the striatum during development. Our study provides data on the developmental mechanism of the claustrum in primates that is fundamental to revealing neuronal circuitry establishment of higher cognitive functions and their pathogenesis in humans.

## Materials and Methods

### Animals

All animal experiments were approved by the Institutional Animal Care & Use Committee (IACUC) at Yale University. Animal breeding and dating of pregnancies have been described in the previous publications (46, 47). Briefly, pregnant macaques were validated by chorionic gonadotropin (MCG) testing and ultrasound, each cesarean section was performed at a pre-determined gestational stage followed by post-operative care. For 5-Chloro-2’-deoxyuridine (CldU) and 5-Iodo-2’-deoxyuridine (IdU) injections, animals received either a single intravenous injection of CldU or IdU for single labeling or one injection of CldU and one injection of IdU at different gestational ages for double labeling. CldU or IdU (30mg/ml) were dissolved in 0.9%NaCl, and filter-sterilized before injection at 50mg/Kg body weight.

Material already publicly available in MacBrain Resource collection 1 and 6 were used extensively for controls and comparisons (e.g., in **Fig. 1** and **Fig. 7, Fig. S1, Fig. S2**) and therefore no animals needed to be sacrificed for that part of the study. See: https://medicine.yale.edu/neuroscience/macbrain/

### Histology

Brains from fetuses and postnatal animals were carefully dissected and fixed in 4% paraformaldehyde for 24 hours before soaking into 10%, 20%, and 30% sucrose solution sequentially for cryoprotection. Brains were then divided into 2 to 4 blocks for each hemisphere. Sections were cut in the coronal plane at 40μm to 50μm with a cryostat microtome. Sections were numbered increasingly from anterior to posterior starting at the position where the claustrum appears.

### Immunohistochemistry

Sections were washed with PBS before being incubated in a blocking solution (5% donkey serum in 0.4% Triton X-100 PBS). Antibodies to NR4A2 (1:200, Abcam, ab41917 RRID:AB_776887), SATB2 (1:200, Abcam, ab51502, RRID:AB_882455), TBR1 (1:500, Abcam, ab31940, RRID: AB_2200219), NeuN (1:500, EMD Millipore, ABN78, RRID: AB_10807945), SOX5 (1:200, Abcam, ab94396, RRID: AB_10859923), CTIP2 (1:200, Abcam, ab18465, RRID: AB_2064130), Parvalbumin (1:1000, Sigma, P3088, RRID: AB_477329) were incubated overnight at 4°C in PBS with 2% donkey serum. After washing three times in PBST, sections were incubated with species-specific secondary antibodies for 2hr at room temperature. Secondary antibodies were obtained from Jackson ImmunoResearch. For single immunofluorescence labeling, either species-antibodies labeled with Alexa 488 or Cy3 were used. For double labeling, Alexa 488 was combined with Cy3 or Cy5. Sections were washed again three times and then mounted with Vectashield Mounting Medium for Fluorescence (Vector Laboratories Inc.).

### Fluorescence imaging

Single channel and multichannel Images were acquired with ZEISS LSM 880 with Airyscan (Carl Zeiss, Germany). 10X and 20X objectives were used for whole slide tile imaging, and 20X and 40X objectives were used to acquire single optical section images for colocalization analysis. Stitching tile images was carried out by ZEN DESK Blue software and images were then exported into tiff files. Some whole slide scan images were acquired with AxioScan7 fluorescence scanner (Carl Zeiss, Germany), 10X and 20X objectives were used.

## Acknowledgments

This study was supported by NIH grants 2R37DA023999-12 (P.R.) and 2R01MH113257-06 (A.D.).

## References

1. F. C. Crick, C. Koch, What is the function of the claustrum? Philos. Trans. R. Soc. Lond. B. Biol. Sci. 360, 1271–9 (2005).

2. J. S. Baizer, C. C. Sherwood, M. Noonan, P. R. Hof, Comparative organization of the claustrum: what does structure tell us about function? Front. Syst. Neurosci. 8, 117 (2014).

3. J. S. Baizer, C. J. Webster, J. F. Baker, The Claustrum in the Squirrel Monkey. Anat. Rec. 303, 1439–1454 (2020).

4. S. LeVay, H. Sherk, The visual claustrum of the cat. II. The visual field map. J. Neurosci. 1, 981–992 (1981).

5. S. LeVay, H. Sherk, The visual claustrum of the cat. I. Structure and connections. J. Neurosci. 1, 956–980 (1981).

6. D. Minciacchi, A. Granato, P. Barbaresi, Organization of claustro-cortical projections to the primary somatosensory area of primates. Brain Res. 553, 309– 312 (1991).

7. Y. Goll, G. Atlan, A. Citri, Attention: The claustrum. Trends Neurosci. 38, 486–495 (2015).

8. J. Kim, C. J. Matney, R. H. Roth, S. P. Brown, Synaptic Organization of the Neuronal Circuits of the Claustrum. J. Neurosci. 36, 773–84 (2016).

9. G. Atlan, A. Terem, N. Peretz-Rivlin, M. Groysman, A. Citri, Mapping synaptic cortico-claustral connectivity in the mouse. J. Comp. Neurol. 525, 1381–1402 (2017).

10. M. G. White, et al., Anterior Cingulate Cortex Input to the Claustrum Is Required for Top-Down Action Control. Cell Rep. 22, 84–95 (2018).

11. G. Atlan, et al., The Claustrum Supports Resilience to Distraction. Curr. Biol. 28, 2752-2762.e7 (2018).

12. L. Kurada, A. Bayat, S. Joshi, M. Z. Koubeissi, The Claustrum in Relation to Seizures and Electrical Stimulation. Front. Neuroanat. 0, 8 (2019).

13. H. Norimoto, et al., A claustrum in reptiles and its role in slow-wave sleep. Nature 578, 413–418 (2020).

14. K. Narikiyo, et al., The claustrum coordinates cortical slow-wave activity. Nat. Neurosci. 23, 741–753 (2020).

15. M. Chevée, E. A. Finkel, S. J. Kim, D. H. O’Connor, S. P. Brown, Neural activity in the mouse claustrum in a cross-modal sensory selection task. Neuron 110, 486–501.e7 (2022).

16. T. R. Vidyasagar, E. Levichkina, An Integrated Neuronal Model of Claustral Function in Timing the Synchrony Between Cortical Areas. Front. Neural Circuits 13, 3 (2019).

17. M. G. White, et al., The Mouse Claustrum Is Required for Optimal Behavioral Performance Under High Cognitive Demand. Biol. Psychiatry 88, 719–726 (2020).

18. H.-G. Bernstein, et al., Bilaterally reduced claustral volumes in schizophrenia and major depressive disorder: a morphometric postmortem study. Eur. Arch. Psychiatry Clin. Neurosci. 266, 25–33 (2016).

19. M. C. Patru, D. H. Reser, A New Perspective on Delusional States -Evidence for Claustrum Involvement. Front. psychiatry 6, 158 (2015).

20. N. G. Cascella, G. J. Gerner, S. C. Fieldstone, A. Sawa, D. J. Schretlen, The insula-claustrum region and delusions in schizophrenia. Schizophr. Res. 133, 77– 81 (2011).

21. D. Binks, C. Watson, L. Puelles, A Re-evaluation of the Anatomy of the Claustrum in Rodents and Primates—Analyzing the Effect of Pallial Expansion. Front. Neuroanat. 13, 34 (2019).

22. L. Puelles, “Development and Evolution of the Claustrum” in The Claustrum: Structural, Functional, and Clinical Neuroscience, (Elsevier Inc., 2014), pp. 119– 176.

23. J. B. Smith, et al., The relationship between the claustrum and endopiriform nucleus: A perspective towards consensus on cross-species homology. J. Comp. Neurol. 527, 476–499 (2019).

24. H. Bruguier, et al., In search of common developmental and evolutionary origin of the claustrum and subplate. J. Comp. Neurol. (2020) https://doi.org/10.1002/cne.24922 (August 1, 2020).

25. P. Rakic, Neurons in rhesus monkey visual cortex: systematic relation between time of origin and eventual disposition. Science 183, 425–427 (1974).

26. K. Y. Kwan, et al., SOX5 postmitotically regulates migration, postmigratory differentiation, and projections of subplate and deep-layer neocortical neurons. Proc. Natl. Acad. Sci. U. S. A. 105, 16021–16026 (2008).

27. Y. Arimatsu, I. Nihonmatsu, Y. Hatanaka, Localization of latexin-immunoreactive neurons in the adult cat cerebral cortex and claustrum/endopiriform formation. Neuroscience 162, 1398–1410 (2009).

28. Y. Hatanaka, et al., Intracortical Regionality Represented by Specific Transcription for a Novel Protein, Latexin. Eur. J. Neurosci. 6, 973–982 (1994).

29. R. Druga, S. Chen, M. Bentivoglio, Parvalbumin and calbindin in the rat claustrum: An immunocytochemical study combined with retrograde tracing from frontoparietal cortex. J. Chem. Neuroanat. 6, 399–406 (1993).

30. B. N. Mathur, The claustrum in review. Front. Syst. Neurosci. 8, 48 (2014).

31. B. Rudy, G. Fishell, S. H. Lee, J. Hjerling-Leffler, Three groups of interneurons account for nearly 100% of neocortical GABAergic neurons. Dev. Neurobiol. 71, 45–61 (2011).

32. I. H. Smart, M. Smart, Growth patterns in the lateral wall of the mouse telencephalon: I. Autoradiographic studies of the histogenesis of the isocortex and adjacent areas. J. Anat. 134, 273–98 (1982).

33. I. H. Smart, M. Smart, The location of nuclei of different labelling intensities in autoradiographs of the anterior forebrain of postnatial mice injected with [3H]thymidine on the eleventh and twelfth days post-conception. J. Anat. 123, 515–51525 (1977).

34. A. Hoerder-Suabedissen, et al., Temporal origin of mouse claustrum and development of its cortical projections. Cereb. Cortex (2022) https://doi.org/10.1093/CERCOR/BHAC318 (October 27, 2022).

35. Z. Chia, G. J. Augustine, G. Silberberg, Synaptic Connectivity between the Cortex and Claustrum Is Organized into Functional Modules. Curr. Biol. 30, 2777–2790.e4 (2020).

36. R. Remedios, N. K. Logothetis, C. Kayser, Unimodal Responses Prevail within the Multisensory Claustrum. J. Neurosci. 30, 12902–12907 (2010).

37. Q. Wang, et al., Organization of the connections between claustrum and cortex in the mouse. J. Comp. Neurol. 525, 1317–1346 (2017).

38. J. A. Gorski, et al., Cortical excitatory neurons and glia, but not GABAergic neurons, are produced in the Emx1-expressing lineage. J. Neurosci. 22, 6309–14 (2002).

39. B. Rudy, G. Fishell, S. H. Lee, J. Hjerling-Leffler, Three groups of interneurons account for nearly 100% of neocortical GABAergic neurons. Dev. Neurobiol. 71, 45–61 (2011).

40. K. Reynhout, J. S. Baizer, Immunoreactivity for calcium-binding proteins in the claustrum of the monkey. Anat. Embryol. (Berl). 199, 75–83 (1999).

41. M. Á. Real, J. C. Dávila, S. Guirado, Expression of calcium-binding proteins in the mouse claustrum. J. Chem. Neuroanat. 25, 151–160 (2003).

42. J.-I. Johnson, B. A. Fenske, A. S. Jaswa, J. A. Morris, Exploitation of puddles for breakthroughs in claustrum research. Front. Syst. Neurosci. 8, 78 (2014).

43. N. Patzke, G. M. Innocenti, P. R. Manger, The claustrum of the ferret: afferent and efferent connections to lower and higher order visual cortical areas. Front. Syst. Neurosci. 8, 31 (2014).

44. J. B. Smith, A. K. Lee, J. Jackson, The claustrum. Curr. Biol. 30, R1401–R1406 (2020).

45. R. C. A. Pearson, P. Brodal, K. C. Gatter, T. P. S. Powell, The organization of the connections between the cortex and the claustrum in the monkey. Brain Res. 234, 435–441 (1982).

46. P. Rakic, Kinetics of proliferation and latency between final cell division and onset of differentiation of cerebellar stellate and basket neurons. J. Comp. Neurol. 147, 523–546 (1973).

47. D. R. Kornack, P. Rakic, Changes in cell-cycle kinetics during the development and evolution of primate neocortex. Proc. Natl. Acad. Sci. U. S. A. 95, 1242–1246 (1998).

