## Supplementary material for "Origin and Development of the Claustrum in Rhesus Macaque"

**This PDF file includes:**

Fig. S1 to S2

Fig. S1

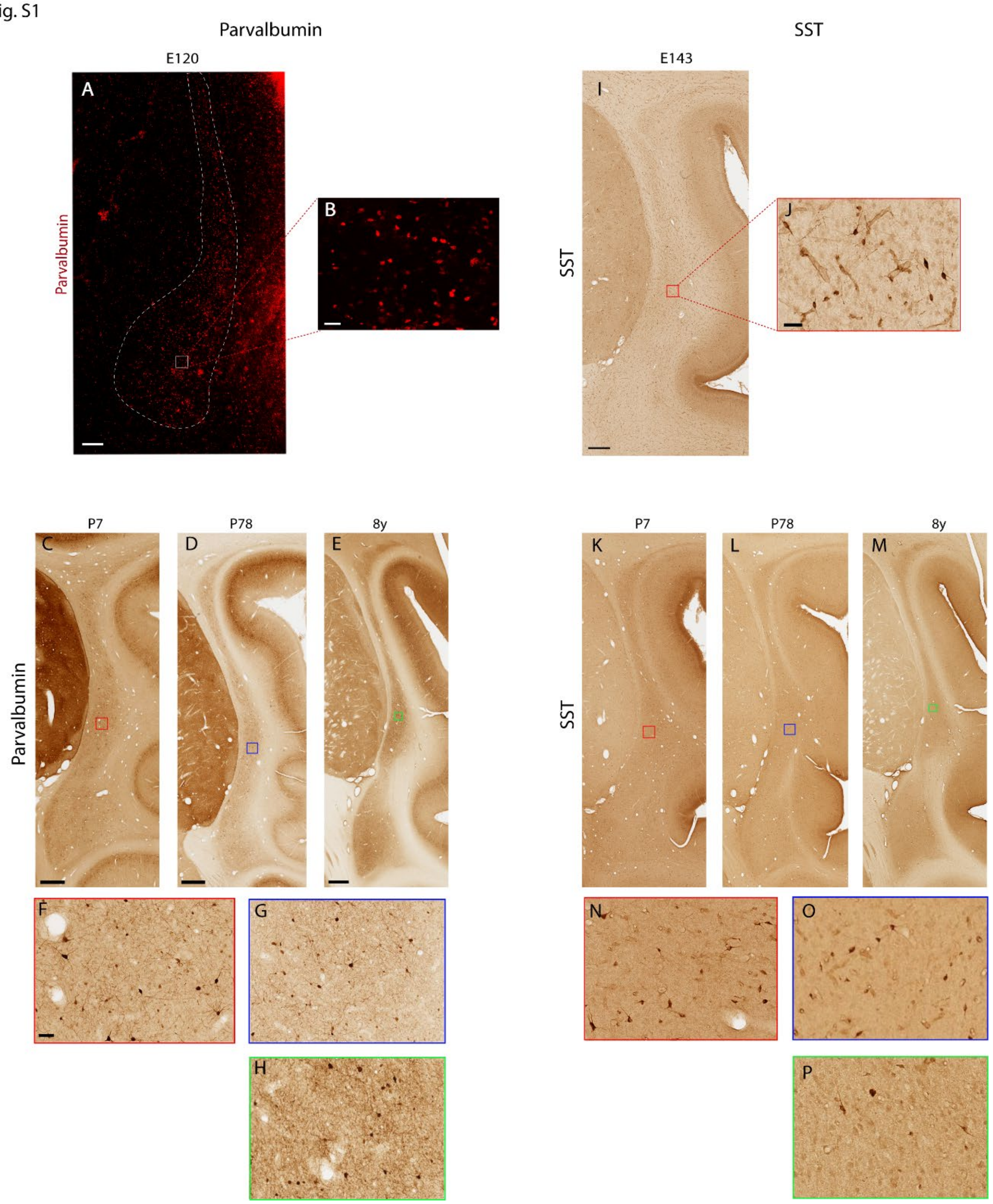


**Fig. S1.** Parvalbumin and SST interneurons in the developing claustrum. (**A-H**) Parvalbumin positive neurons are detected abundantly from E120 to adult. (**B**, **F**, **G**) are close-up images from the framed area in (**A**, **C**, **D**, **E**) respectively. SST positive neurons are detected from E143 to adult. (**J**, **N**, **O**, **P**) are close-up images from the framed area in (**I**, **K**, **L**, **M**) respectively. Scale bar: A: 250μm; B, J: 40μm; C, I, K: 1mm; D, L: 1mm; E, M: 1mm; F-H, N-P: 40μm. Images are from MacBrainResource Collection 6, Brain 64 (P7), Brain 65 (P78), Brain 72 (8y).

Fig. S2

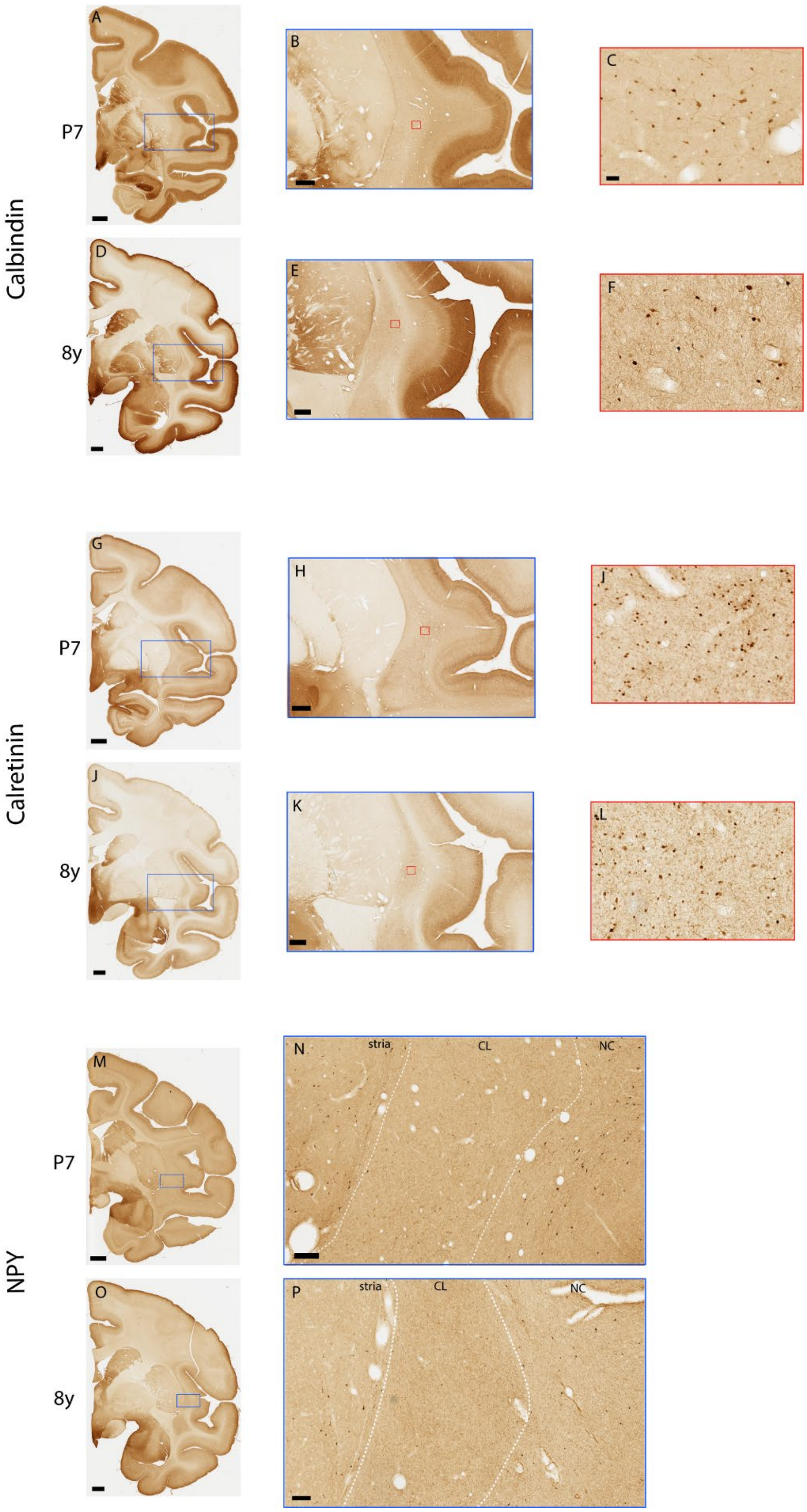

Calbindin
P7
8y
A
B
C
D
E
F
Calretinin
G
H
I
J
K
L
NPY
M
N
O
P
stria
CL
NC

**Fig. S2.** Calbindin, Calretinin and NPY positive interneurons in the developing claustrum. Calbindin (**A**-**F**) and Calretinin (**G**-**L**) positive interneurons are present in the claustrum of postnatal and adult brains. (**B**, **E**) are enlarged images from blue framed areas in (**A**) and (**D**). (**C**, **F**) are close up images from red framed areas in (**B**) and (**E**). (**H**, **K**) are enlarged images from blue framed areas in (**G**) and (**J**). (**I**, **L**) are close-up images from red framed areas in (**H**) and (**K**). However, NPY expression is low in the claustrum (**M**-**P**). (**N**, **P**) are close-up images from (**M**) and (**O**). In contrast, NPY positive neurons are present in striatum (stria) and neocortex (NC). White dashed lines in (**N**) and (**P**) mark the borderline of the claustrum (CL). Scale bar: A, G, M: 2mm; B, H: 500μm. D, J, O: 2mm; E, K, N, P: 500μm; C, F, **I**, L: 70μm. Images are from MacBrainResource Collection 6, Brain 64 (P7), Brain 72 (8y).
